# Transcriptomics provides a genetic signature of vineyard site with insight into vintage-independent regional wine characteristics

**DOI:** 10.1101/2021.01.07.425830

**Authors:** Taylor Reiter, Rachel Montpetit, Shelby Byer, Isadora Frias, Esmeralda Leon, Robert Viano, Michael Mcloughlin, Thomas Halligan, Desmon Hernandez, Rosa Figueroa-Balderas, Dario Cantu, Kerri Steenwerth, Ron Runnebaum, Ben Montpetit

## Abstract

In wine fermentations, the metabolic activity of both *Saccharomyces cerevisiae* and non-*Saccharomyces* organisms impact wine chemistry. Ribosomal DNA amplicon sequencing of grape musts has demonstrated that microorganisms occur non-randomly and are associated with the vineyard of origin, suggesting a role for the vineyard, grape, and wine microbiome in shaping wine fermentation outcomes. We used ribosomal DNA amplicon sequencing of grape must and RNA sequencing of primary fermentations to profile fermentations from 15 vineyards in California and Oregon across two vintages. We find that the relative abundance of fungal organisms detected by ribosomal DNA amplicon sequencing did not correlate with transcript abundance from those organisms within the RNA sequencing data, suggesting that the majority of the fungi detected in must by ribosomal DNA amplicon sequencing are not active during these inoculated fermentations. Additionally, we detect genetic signatures of vineyard site and region during fermentation that are predictive for each vineyard site, identifying nitrogen, sulfur, and thiamine metabolism as important factors for distinguishing vineyard site and region.

**Importance:** The wine industry generates billions of dollars of revenue annually, and economic productivity is in part associated with regional distinctiveness of wine sensory attributes. Microorganisms associated with grapes and wineries are influenced by region of origin, and given that some microorganisms play a role in fermentation, it is thought that microbes may contribute to the regional distinctiveness of wine. We show that while the presence of microbial DNA is associated with wine region and vineyard site, the presence of microbial DNA is not associated with gene expression of those microorganisms during fermentation. We further show that detected gene expression signatures associated with wine region and vineyard site provide a means to address differences in fermentations that may drive regional distinctiveness.

## Introduction

During vinification, grape musts are transformed to wine through microbial metabolism, including fermentation of grape sugars into alcohols. In both inoculated and spontaneous fermentations, *Saccharomyces cerevisiae* often becomes the dominant fermentative organism due to a milieu of adaptations that support the rapid consumption of sugars and production of ethanol (1). However, complex microbial communities consisting of other eukaryotic microorganisms and bacteria are present, active, and make significant contributions to the wine making process and final product (2–6). Referred to collectively as non-*Saccharomyces* organisms, these microbes often originate from the vineyard or the winery itself (7, 8). In recognition of the important role these microbes have in wine making, select non-*Saccharomyces* yeasts are increasingly inoculated into commercial fermentations to impart beneficial properties (e.g. bio-protection, lower ethanol, or distinct sensory characteristics (9)). Grape must treatment with sulfur dioxide (SO_2_) is also commonly used to control microbial populations, including spoilage organisms, but many microorganisms survive SO_2_ treatment and contribute to fermentation outcomes (6, 10, 11).

The persistence of vineyard and winery derived microorganisms throughout the winemaking process, as well as the potential for these organisms to influence grape berry development prior to harvest, has led to the idea that select microorganisms unique to a region or vineyard may contribute to region-specific wine characteristics (12, 13). In support of a role of microbial biogeography in regional wine characteristics, microorganisms in vineyards, wineries, and grape musts are known to be associated with their region of origin (4, 7, 8, 14–21). Moreover, the abundance of some organisms in grape must correlates with metabolite concentrations in finished wine, further associating microbial biogeography to fermentation outcomes and wine quality (15, 22). Still, relatively little is known about the influence of non-*Saccharomyces* microorganisms on wine fermentation outcomes, but an increasing number of studies are tackling this complex problem (23, 24). Recent studies have documented increased glycerol accumulation and aroma profiles using sequential- or co-inoculation of *S. cerevisiae* with a single non-*Saccharomyces* yeast species under enological conditions (25–34). While outcomes are diverse, which may be expected given the variety of starting must and culture conditions used across studies, many report consistent alterations in wine such as a higher glycerol content from fermentations inoculated with *S. cerevisiae* and *Starmerella bacillaris* (29, 30, 34).

How these altered fermentation outcomes occur remains a difficult question to address, as a given outcome may be the direct result of metabolism by the non-*Saccharomyces* organism, or the result of the organism altering *S. cerevisiae* metabolism via direct or indirect interactions (35–37). In support of the latter, the presence of non-*Saccharomyces* organisms has been shown to increase the rate and diversity of resource uptake by *S. cerevisiae* in early fermentation (36–38). In controlled steady-state bioreactor fermentations, the presence of *Lachancea thermotolerans* was found to increase the expression of *S. cerevisiae* genes important for iron and copper acquisition (39). Such interactions are not limited to fungi—lactic acid bacteria can induce epigenetic changes (e.g. *[GAR+]* prion) in *S. cerevisiae* that alter glucose metabolism (40–42). Such abilities of non-*Saccharomyces* organisms to impact *S. cerevisiae* metabolism and fermentation outcomes raises the question of whether microbial biogeography of vineyard sites persists in fermentations, thereby influencing wine outcomes in a site-specific manner. In addition, microbial diversity changes as primary fermentation progresses and *S. cerevisiae* becomes dominant (43), suggesting a changing microbial community could feedback to impact fermentation progression in multiple distinct ways. Currently, we know relatively little about these inter-species interactions and how this influences *S. cerevisiae*, which as a field must be addressed if we are to understand how microbial community dynamics impact wine fermentation outcomes.

Past inquiries into the microbial communities of grape must and wine related to regional distinctiveness have focused on assaying the presence of specific microbes based on ribosomal DNA amplicon sequencing (4, 8, 14–20, 44). DNA sequencing has the advantages of capturing both metabolically active and inactive organisms, due to the relative stability of the DNA molecule, offering evidence of a rich history of the microbial community prior to sampling. Ribosomal DNA amplicon data further provides a measure of what microbes may be active at the time of sampling or may become active in the future. While microbiome DNA sequencing of grape musts supports regionally distinct microbial signatures, what microbes are metabolically contributing to fermentation outcomes remains largely unknown. This information is critical when considering the possibility that a particular microbe influences a wine fermentation outcome via metabolism or inter-species interactions.

One measure of metabolic activity that is relatively accessible and can be applied at scale to address this issue is the measurement of gene expression in both *S. cerevisiae* and other non-*Saccharomyces* organisms. An interrogation of the genes that are “on” at a given time using RNA sequencing provides important information about the activities an organism may be performing. In addition, the RNA molecule assessed by transcriptomics is constantly turned over within cells and is relatively unstable compared to DNA, which we propose makes transcriptomics a good indicator of microbial activity at the time of sampling. For example, early in fermentation *S. cerevisiae* turns on genes required for glucose metabolism and represses expression of genes needed for the metabolism of other carbon sources; a pattern that reverses towards the end of fermentation when glucose is depleted and *S. cerevisiae* must find alternative energy sources (45). These patterns of gene expression are easily observed using transcriptomics (45, 46), which is increasingly being applied to understanding wine fermentation outcomes (36–39, 47).

Here, we characterize microbial populations present in Pinot noir musts from California and Oregon in multiple vintages using both ribosomal DNA amplicon data from grape must samples and gene expression data from multiple fermentation timepoints. We demonstrate that genetic signatures (i.e., DNA and RNA profiles) of vineyard site and region are captured by these data, with total precipitation during growing season being one vineyard-associated factor identified to correlate with site-specific genetic signatures. While DNA profiles reliably predict both vineyard site and region, these profiles did not correlate with the RNA profiles of the primary fermentations. This finding suggests other characteristics influence site-specific gene expression signatures more than the grape must microbiome as measured by ribosomal DNA amplicon sequencing. Importantly, a comparison of DNA sequencing and gene expression data indicates that the majority of organisms detected by ribosomal DNA sequencing lack detectable gene expression during the primary fermentation, thus limiting the likelihood that many of these organisms significantly impact fermentation outcomes during the primary stage of fermentation. Finally, using *S. cerevisiae* gene expression patterns and the associated functions of the genes identified, we are able to identify candidate factors that contribute to vineyard specific fermentation outcomes and wine sensory characteristics.

## Results and Discussion

To investigate associations between grape must microbial communities and regional distinctiveness of resulting wines, we performed standardized fermentations of Pinot noir grapes from 15 vineyard sites in California and Oregon across multiple vintages (**Figure S1A**). In 2016, 2017, and 2019, we performed four inoculated fermentations per vineyard site using the wine yeast RC212, taking microbiome samples for DNA isolation and ribosomal DNA amplicon sequencing prior to inoculation. In the 2017 and 2019 vintages, we further profiled two primary fermentations from each site using RNA sequencing approaches to perform gene expression analyses at multiple fermentation timepoints. We performed all grape processing and temperature-controlled fermentations at the UC Davis Teaching & Research Winery to standardize vinification and minimize contributions from other factors (winery and winemaker) to the microbiome and transcriptome (48–50).

### DNA abundance by ribosomal amplicon sequencing is a poor predictor of detectable gene expression during fermentation

Using ribosomal DNA amplicon sequencing of bacteria and fungi, we detected 3254 distinct bacterial sequences and 2452 distinct fungal sequences in grape must (**Figure 1A and 1B**), with a greater mean species diversity per vineyard site for bacteria than for fungi (**Figure S1B**). However, the core microbiome - i.e., the species present in 90% of all grape musts across all vintages with at least 1% abundance - was larger for fungi than bacteria. The core microbiome consisted of 11 bacterial variants classified to nine taxonomic ranks and 19 fungal variants classified to 10 taxonomic ranks. All bacteria in the core microbiome belonged to the phylum *Proteobacteria* and were dominated by the genus *Tatumella*. (**Figure S2**). *Tatumella* has previously been identified as a dominant genera in other red wine fermentations where it correlated with total acid (by titration) in grape must (51), however these associations have not been experimentally validated. Three of the most abundant bacterial sequence variants identify to the acetic acid producing genus *Gluconobacter* (**Figure S2**). *Gluconobacter* is one of three genera of acetic acid bacteria associated with wine spoilage and the only genus we identify among dominant organisms (**Figure S2**) (52). *Gluconobacter* are primarily active in grape must as the wine environment restricts growth of organisms in this genus (52). Fungi in the core microbiome belong to a single phylum, *Ascomycota*, with all fermentations dominated by the genus *Hanseniaspora*, in particular *Hanseniaspora uvarum*. *H. uvarum* cannot complete alcoholic fermentation alone, but participates in and can alter the quality outcomes of wine fermentations (53). We also identified the fungal genus *Botrytis* among dominant organisms (**Figure S2**), although we lacked the ability to resolve whether the particular variant we detected belongs to the spoilage organism *Botrytis cinerea* or another species in the *Botrytis* genus. Through this work, we have extended microbiome must sequencing to include the 2019 vintage, with results largely matching findings from previous vintages across these same vineyard sites (50). The observed microbial community composition was consistent with organisms previously shown to be present at the initial stages of the wine making process (4, 15–17, 51).

**Figure 1:**
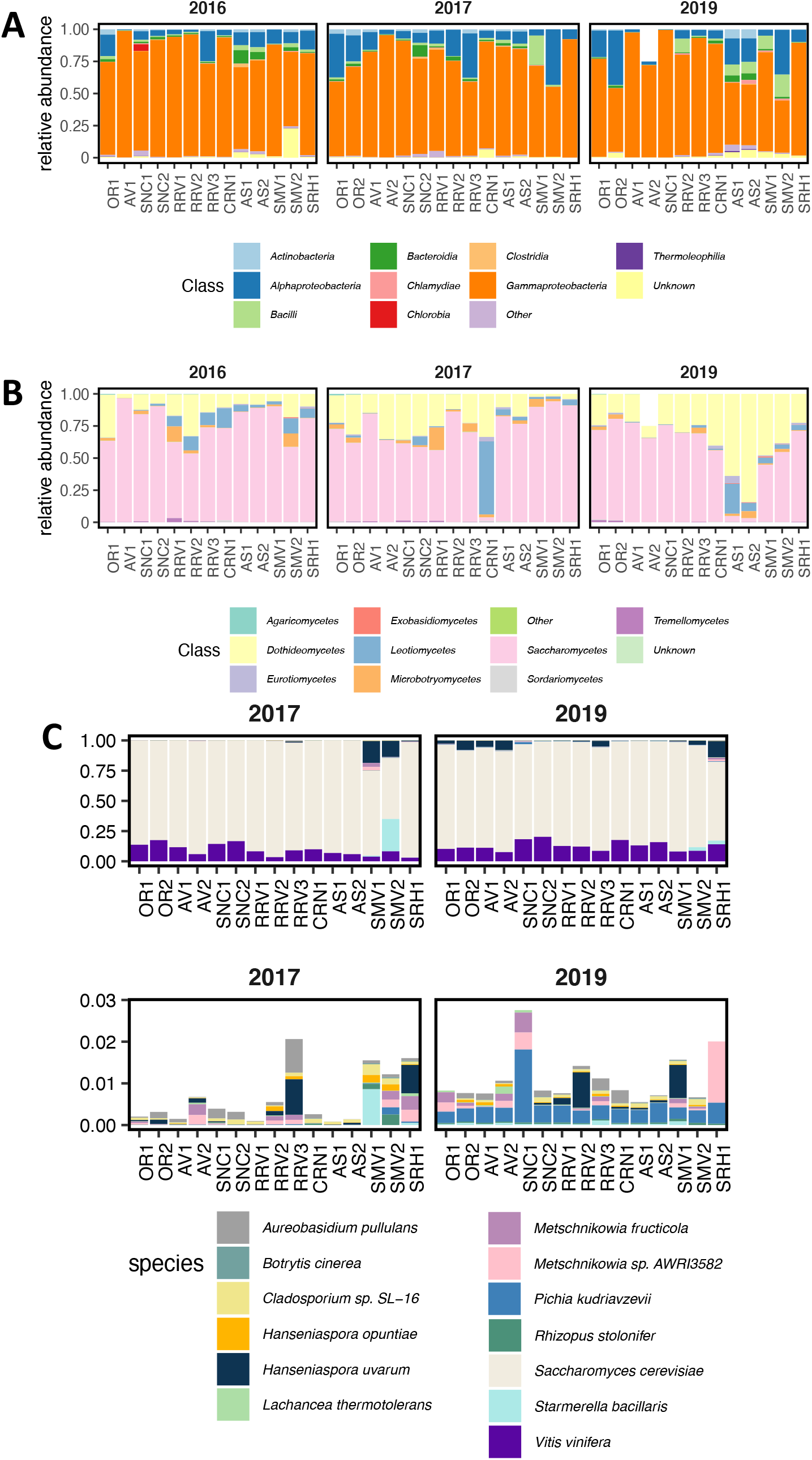
Microbial diversity in grape must and fermentation microbiomes from different vineyard sites. **A, B)** Relative abundance of taxonomic ranks in ribosomal DNA amplicon sequencing data capturing **A** Bacteria and **B** Fungi. Samples taken from fermentations from the same vineyard site and vintage are combined together and reflect relative abundance of organisms from four fermentation tanks. Only three tanks were fermented for AV2 in 2019 due to a smaller harvest. **C)** Relative abundance of all genes expressed by a detected organism during fermentation from the 2017 and 2019 vintages. The top plots show all organisms and bottom plots display only those organisms that account for less than 3% of mapped reads in each sample. Only organisms present in more than one fermentation are plotted.

Ribosomal DNA amplicon sequencing of grape must is expected to capture cells that are metabolically active, inactive, or dead due to the stability of the DNA molecule. In contrast, gene expression profiling via RNA sequencing is expected to be biased towards living cells. Moreover, the identity of the gene transcripts present at the time of sampling further provides information about what metabolic activities the cell may be performing. Using 3’ Tag RNA sequencing (3’ Tag-seq), we profiled eukaryotic organisms during fermentation using samples taken at multiple fermentation timepoints (i.e., 16, 64, and 112 hours after inoculation in 2017 and 2019, plus 2 and 6 hours post-inoculation in 2019). While traditional RNA-sequencing produces sequencing reads from an entire transcript, 3’ Tag-seq produces one molecule per transcript by sequencing approximately 100 base pairs upstream of the 3’-end of a sequence (54). This sequencing chemistry requires a poly(A) tail, limiting the sequenced fraction of the transcriptome almost entirely to polyadenylated eukaryotic mRNAs.

From the resulting 3’ Tag-seq data, we observed that relatively few eukaryotic microbes were detected during these Pinot noir fermentations (**Figure 1C**). Considering all 15 sites together, only 18 eukaryotic species were detected. Further reflecting this finding, *S. cerevisiae* transcripts accounted for the majority of sequences across all fermentations at all time points. To assess whether non-inoculated *S. cerevisiae* strains were responsible for some fraction of sequence reads, we compared each transcriptome against all annotated *S. cerevisiae* genomes in GenBank, as well as a genome assembly of *S. cerevisiae* RC212. While non-RC212 *S. cerevisiae* strains were detectable in every fermentation, this fraction accounted for less than 1% of uniquely identifiable sequences. This demonstrates that the inoculated RC212 strain dominated fermentations at all sampled time points. Interestingly, we also identified *Vitis vinifera* transcripts in all samples (**Figure 1C**). The presence of *V. vinifera* transcripts suggests intact grape cells persist throughout fermentation.

In comparing specific organisms detected via DNA sequencing and 3’ Tag-seq RNA sequencing, we see that only four (*Aureobasidium pullulans, Hanseniaspora uvarum, Hanseniaspora vineae*, and *S. cerevisiae*) of 397 distinct fungal species definitively identified by ribosomal DNA profiling were detected using gene expression data. This was unchanged in the 2019 transcriptome profiling samples taken at 2 and 6 hours after inoculation, suggesting that organisms detected by amplicon sequencing had lost activity prior to or concurrent with inoculation, well before *S. cerevisiae* would begin to produce inhibitory concentrations of ethanol. Of the four detected organisms by 3’ Tag-seq, ribosomal DNA amplicon sequencing data indicated that *H. uvarum* was most abundant in musts prior to inoculation and was detected in every vineyard site (**Figure 2A**). Still, the relative abundance of *H. uvarum* in grape must from ribosomal DNA amplicon sequencing was only weakly correlated with relative abundance of RNA during fermentation (R^2^ = 0.14, p < 0.01). Importantly, while these values are weakly correlated, *H. uvarum* had almost no detectable gene expression in fermentations from many sites where it dominated the DNA profile of the grape must (**Figure 2B**). Finally, even when we performed this analysis using samples from the first hours of fermentation after inoculation, relative abundance of *H. uvarum* DNA in grape musts remained weakly correlated with relative abundance of RNA (two hours: R^2^ = 0.21, p < 0.05, six hours: R^2^ = 0.28, p < 0.01). In the case of *A. pullulans*, DNA in grape must is not correlated with gene expression during fermentation (two hours: R^2^ = −0.03, p = 0.60, six hours: R^2^ = −0.025, p = 0.53). These results indicate that most identified eukaryotic microorganisms in grape must by DNA profiling likely have little metabolic activity in inoculated fermentations even when the organisms are detected at high abundance and are detectable via both sequencing methods.

**Figure 2:**
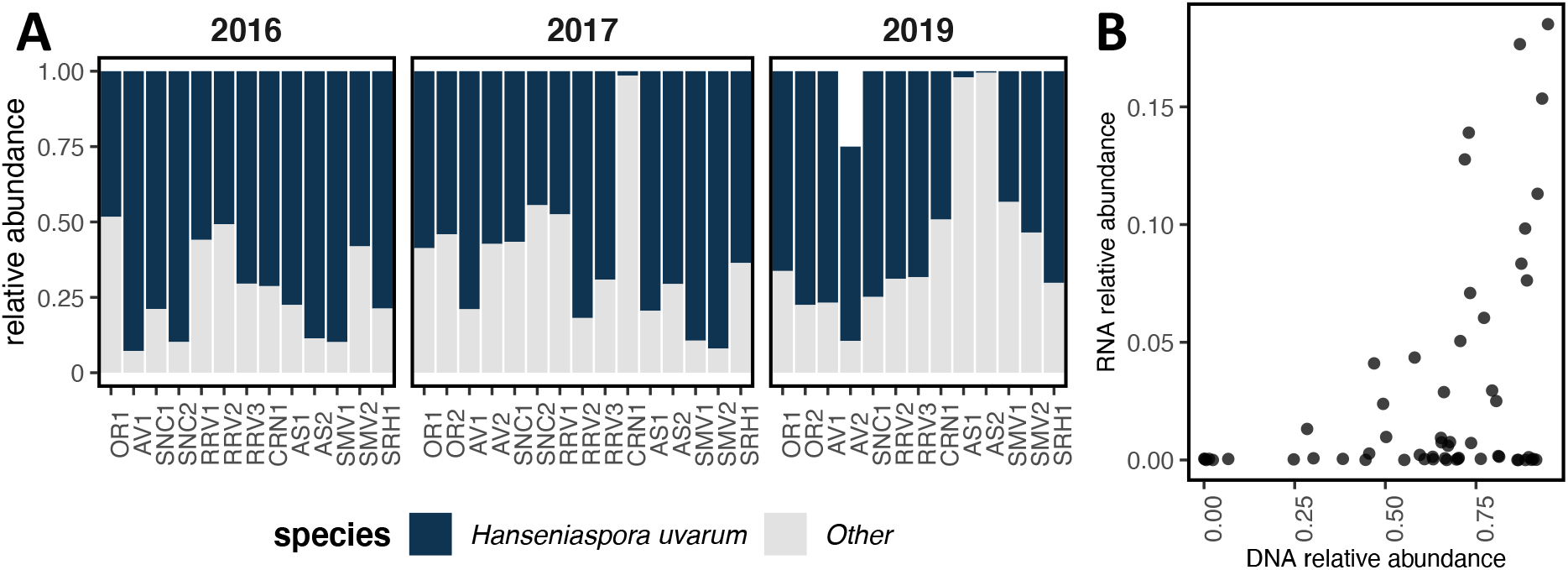
*H. uvarum* ribosomal DNA amplicon sequencing data does not strongly correlate with relative abundance in RNA sequencing data. **A** Bar chart of relative abundance of *H. uvarum* compared to other non-*Saccharomyces* species across fermentations from each site based on amplicon sequencing data of ribosomal DNA. **B** Scatter plots relative abundance of *H. uvarum* as determined by amplicon sequencing of ribosomal DNA (x-axis) vs. RNA sequencing (y-axis).

Given these findings, it is important to consider if a lack of detectable gene expression for non-*Saccharomyces* fungal species could be reflective of some other issue that is technical or biological in nature. We consider this highly unlikely for two reasons. First, both DNA and RNA sequencing require similar protocols for extraction of nucleic acids from cells that should perform approximately equally across samples. Second, RNA sequencing relies on a highly conserved biological processes (mRNA polyadenylation), hence while we could envision RNA sequencing failing for one or a few organisms, it should not fail across many fermentations for the large majority of organisms seen in this work. Moreover, of the 16 non-*Saccharomyces* fungi detected via RNA-sequencing, eight of these organisms were not detected at the genus level by DNA profiling (*Cladosporium sp SL-16, Lachancea thermotolerans, Metschnikowia fructicola, Metschnikowia sp. AWRI3582, Pichia kudriavzevii, Preussia sp. BSL10, Rhizopus stolonifer, Starmerella bacillaris*). This suggests that transcriptomic profiling is a sensitive assay able to detect organisms present in a population that are missed by ribosomal DNA amplicon sequencing, which is likely due to an inability to resolve genus or species using ribosomal DNA sequences.

Notably, some of the organisms detected by RNA sequencing have the ability to influence fermentation outcomes: in mixed fermentations with *S. cerevisiae, S. bacillaris* has been shown to lower the final ethanol concentration and increase the concentration of glycerol (55), while *M. fructicola* increased the concentration of esters and terpenes (56). Therefore, the detection of these organisms by RNA sequencing provides valuable information with respect to the potentially active microbial population in these fermentations. Our findings align well with another recent report that showed an RNA-based sequencing strategy is a highly sensitive alternative to amplicon sequencing (57). As such, it may be appropriate to use RNA sequencing as a general method to capture the metabolically active microbial community during wine fermentation, especially when drawing a connection between the wine microbiome and fermentation outcomes.

### Genetic signatures differentiate vineyard site, region, and vintage

The region and site from which grapes are harvested can have an important influence over the character of a resulting wine based by on a variety of factors (e.g., climate, soil type, grape associated microbes). As such, we considered if the data generated using DNA and RNA sequencing strategies during these Pinot noir fermentations is reflective of vineyard site through the generation of unique genetic signatures. To investigate this concept, we grouped DNA and RNA sequencing samples by vineyard site, region, and vintage to see if there were detectable differences among these groups. Using analysis of similarities (ANOSIM; see methods), we determined that all three factors explain differences among groups of samples, with vineyard site explaining the most group similarity (**Figure 3A-D**). This supports the idea that fermentations have a detectable genetic signature that is reflective of vineyard site.

**Figure 3:**
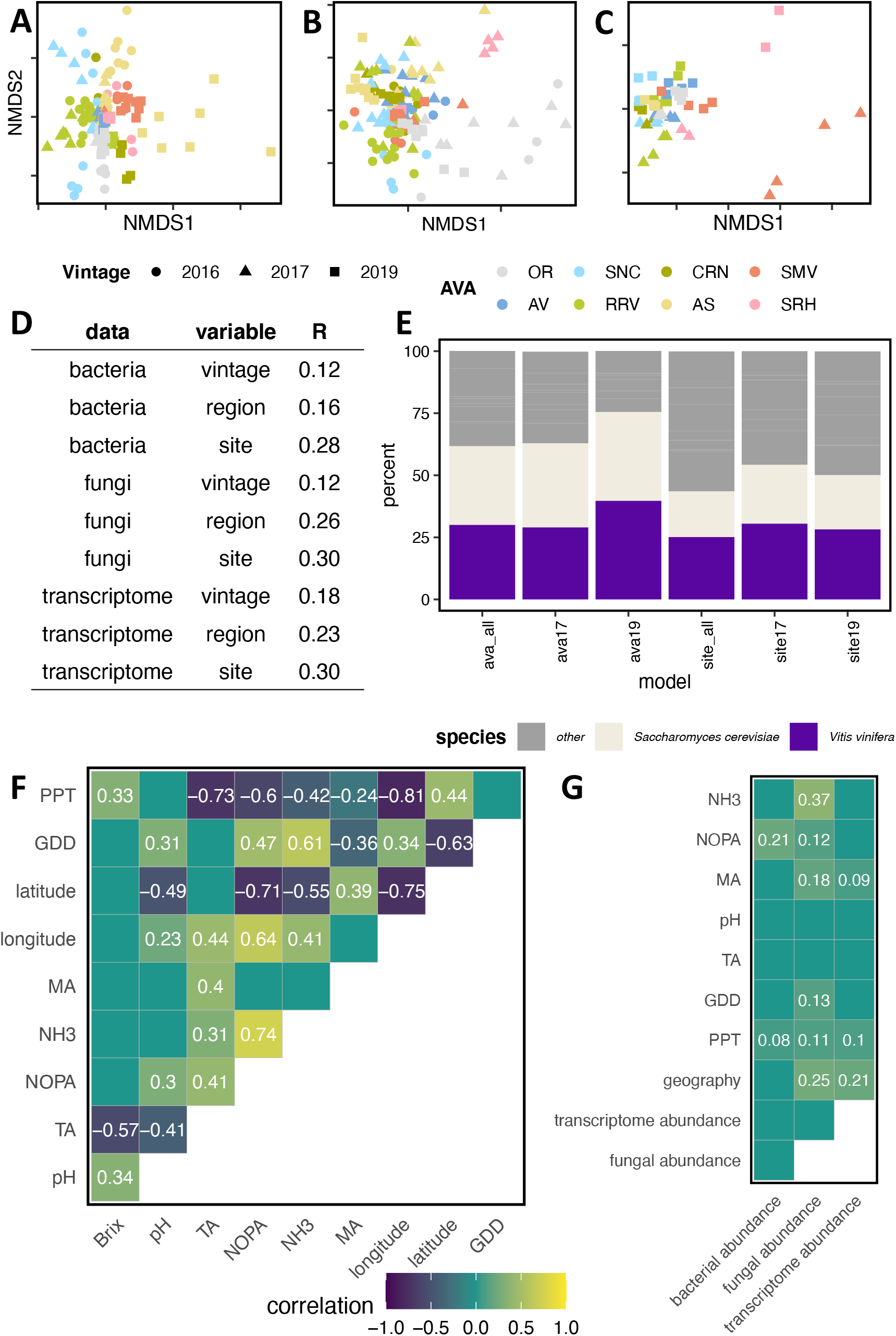
Genetic profiles correlate with vineyard, region, and vintage as well as some vineyard site and initial grape must characteristics. **A-C)** Non-metric Multi-dimensional Scaling plots of Aitchinson dissimilarity of **A** bacterial communities, **B** fungal communities, and **C** and transcriptomes across vintages. The closer two points are on the graph, the more similar their genetic profiles are. **D)** Vineyard site, region, and vintage account for genetic diversity patterns in Analysis of Similarity (ANOSIM). R values represent strength of association, with higher R values indicating stronger grouping according to the parameter. All values are significant (p < 0.001). **E)** Percent of accuracy attributable to different organisms in random forests models. A higher percentage of variable importance was attributable to *S. cerevisiae* and *V. vinifera* in models that predicted region than vineyard site. **F, G)** Correlograms representing similarities between fermentation metrics. **F** Grape must chemical parameters and vineyard site characteristics were correlated in the 2017 and 2019 vintages. Squares are labelled with correlation values from Pearson’s correlation. Only comparisons with p < 0.05 are displayed. **G** Bacterial, fungal, and transcriptome profiles correlated with some vineyard site and grape must chemical characteristics. Squares are labelled with correlation values from Mantel tests. Only comparisons with an FDR < 0.1 are displayed. PPT: precipitation, GDD: growing degree days, MA: malic acid, TA: titratable acidity.

To understand which specific organisms and genes contribute to the genetic signatures of both vineyard site and region, we built machine learning classification models using random forests. These models weight the contribution of each feature to predictive accuracy of the model, enabling robust identification of specific genes or organisms that differentiate vineyard sites or regions among fermentations. When we used data from all vintages in model training and testing to predict region, we achieved 87%-95% accuracy (**Table S1-S3**; **Figure S3–S4**). When we instead used data from one vintage in model training and testing to predict region, accuracy dropped across all models, but ranged from 57%-75% (**Table S1-S3**; **Figure S3–S4**). This suggests that models built with fermentations from all vintages better capture cross-vintage similarities as these models select predictive variables that are consistent across the vintages studied. However, the accuracy of these models may decrease if the same set of predictive variables is not consistent in future vintages. Conversely, the accuracy of a model built from a single vintage and trained on a separate vintage will likely remain consistent across many vintages. From this, we assumed that models trained using data from a single vintage better reflected model accuracy, but that models trained using data from all vintages better reflected cross-vintage similarities. As we aimed to identify vintage-independent factors, we analyzed cross-vintage models moving forward.

When we used the same data to generate vineyard-specific models, predictive accuracy was on average 21.4% less than region-specific models (**Table S1**). However, it is important to note that this decrease in accuracy was driven by within-region misclassification for vineyards in Willamette Valley (31 km separation), Santa Maria Valley (5 km separation), and Arroyo Seco (1 km separation) American Viticultural Areas (AVA) (**Figure S5**). The same misclassifications persisted across many models, highlighting potential within-region similarity that contributes to genetic signatures, which fits well with the concept of AVA and region-associated wine characteristics.

Across models, we were surprised to find that bacterial models outperformed or performed as well as fungal models for classification of site and region, as bacterial must samples added the least predictive power in previous models for region prediction (14), including for Pinot noir grapes grown in Australia (8). Bacterial must samples have been shown to be predictive of region in Californian Chardonnay, but not Californian Cabernet Sauvignon (14), suggesting a possible cultivar-specific effect. In previous inquiries, samples were processed in vineyard-specific wineries, providing another variable that could potentially alter the measured microbiomes and the contributions attributed to bacteria and fungi.

Given that random forests models estimate the importance of each gene in determining vineyard or region classification, we further used the gene expression models to gain insight into biological differences between vineyard sites and regions. For this, we calculated the percent of total importance attributable to each gene from each eukaryotic organism detected (**Table S2**). Vineyard-specific models weighted non-*Saccharomyces* yeast genes as a whole as most important for predictive accuracy (**Figure 3E, Figure S6**). In particular, genes from *S. bacillaris, M. fructicola, Metschnikowia sp. AWR13582*, and *L. thermotolerans* were important for vineyard site classification. The ability of non-*Saccharomyces* gene expression to distinguish site is likely related to the unique combination of non-*Saccharomyces* organisms present in each fermentation, which results in these organisms having strong predictive power when detected. In contrast, regional models weighted *S. cerevisiae* and *V. vinifera* genes as higher importance (**Figure 3E, Figure S6**). We expect that this may result from changes in *V. vinifera* gene expression across more diverse geographical environments, which leads to differences in the grape must and associated fermentations as detected by *S. cerevisiae* gene expression.

To more directly address how environmental factors and grape must chemistry correlate with genetic signatures, we correlated initial must chemical parameters (pH, titratable acidity, malic acid, NOPA, and NH_3_) and vineyard site characteristics (total precipitation, growing degree days, and geographic distance between sites) with DNA and RNA profiles using the Mantel test (see methods). From these analyses, we found geographic distance between vineyards correlated with precipitation and growing degree days, indicating that sites that are geographically closer experience more similar weather patterns, as would be expected (**Figure 3F**). Amongst the factors tested, only precipitation correlated with all genetic profiles (**Figure 3G**). Similar to geographic distance, initial chemical profiles of vineyard sites were more similar when sites are geographically closer. However, we found surprisingly few correlates between genetic profiles and initial grape must conditions (**Figure 3G**). While fungal profiles correlate with initial malic acid, NOPA, and NH_3_ and bacterial profiles correlate with initial NOPA, gene expression profiles only correlate with initial malic acid levels. The finding that gene expression profiles do not correlate with initial nitrogen concentration, even though nitrogen availability is central to yeast growth and linked to the expression of hundreds of genes (45), may reflect nitrogen additions at ~24 h after inoculation during winemaking so that all fermentations had a minimum of 250 mg/L (see methods). Overall, the poor correlation between gene expression patterns and the factors tested suggest that other unmeasured factors drive gene expression distinctiveness in these fermentations. This raises a clear need for future work that measures many factors within vineyards and fermentations to define the organism-environment interactions responsible for driving gene expression and cellular activities of *S. cerevisiae* and other microbial organisms.

### *S. cerevisiae* gene expression provides insight into vineyard site and region features

*S. cerevisiae* is likely the best understood eukaryote based on the use of this organism as a model system for biology, which has provided a rich set of genomic resources and databases (58). As such, *S. cerevisiae* gene expression can be used as a biosensor to provide insight into the fermentation environment based on activities yeast perform. The utility of this data is furthered by the fact that *S. cerevisiae* gene functions are well studied in the context of wine production, *S. cerevisiae* is ubiquitous across all fermentations, and the transcriptomics data is dominated by reads from *S. cerevisiae* (e.g., data completeness). Consequently, given the data above suggesting unknown factors are directing fermentation outcomes, we queried the *S. cerevisiae* gene expression data to assess what genes were important for predicting region and vineyard site to infer what may be unique about musts produced by grapes from each vineyard site or region. Notably, random forests models are non-deterministic, meaning the each time a model is built the specific genes important for predictive accuracy of that model may change, especially for genes with correlated gene expression values (59). Therefore, we first built 100 random forests models for the prediction of region and vineyard site and investigated the genes that were shared across the majority models (**Table S4**). As discussed above, less than 1% of transcripts in any fermentation were expressed by non-RC212 *S. cerevisiae* and thus the genetic signatures we identified are likely specific to this strain.

From this analysis, important predictors of both site and region included flavor-associated genes involved in the formation of higher alcohols and volatile fatty acids through the Ehrlich pathway. Each site-specific and region-specific model included an average of 16 (site SD = 2.9, region SD = 2.4) genes associated with flavor development in wine (**Table S5**). These genes were mostly associated with the Ehrlich pathway (site mean = 8.1 genes, SD = 2; region mean = 9 genes, SD = 1.7) and with volatile sulfur formation (site mean = 6.3 genes, SD = 1.6; region mean = 5.1 genes, SD = 1.4). Given that genes in these pathways were detectable as indicators of both region and site, site-variable expression of these genes could contribute to region- and vineyard-specific wine flavor profiles detected in wines from these vineyards in previous vintages (48). At this time, it remains unknown what factors cause these flavor-associated genes to differ between fermentations.

In addition to flavor-associated genes, many *S. cerevisiae* genes that were important for predicting vineyard site and region are members of the Com2 regulon (**Table S4**). Expression of genes within the Com2 regulon is protective against SO_2_ stress (60). We treated all fermentations with an equal dose of SO_2_ at the beginning of vinification; however, variable application of sulfur-containing fungicides in the vineyard may lead to disparate SO_2_ stress during fermentation underlying the signatures of site and region that we observe. Wine strains of *S. cerevisiae* are more tolerant of SO_2_ than many non-*Saccharomyces* species, but SO_2_ exposure can cause inhibition of key metabolic enzymes like alcohol dehydrogenase, as well as other processes through cleavage of disulfide bonds (61, 62). Of the 511 genes dependent on Com2 activation during SO_2_ stress (60), an average of 105 genes (SD = 12.7) were important for differentiating site in our predictive models, while 101 genes (SD = 11.6) were important for predicting vineyard region. Within these gene lists are genes involved in the efflux of sulfite and bisulfite, sulfate assimilation, sulfate assimilation, biosynthesis of methionine, cysteine, arginine, and lysine, and biosynthesis of the sulfur-containing vitamin biotin (**Table S6**). These pathways, and their site-specific signatures, are potential areas of future study given that sulfur metabolism can have a profound impact on the sensory attributes of wine (63). In addition, while the molecular form of SO_2_ causes *S. cerevisiae* stress and inactivation of wine spoilage microbes (11, 60), this form is in equilibria with the bisulfite form (HSO_3_^-^) and this ratio is dependent on wine pH (64). The bisulfite form interacts with anthocyanins and can cause color bleaching (64). This suggests that the SO_2_ stress response is a factor that would need to be considered in the context of pH and other aspects of SO_2_ wine chemistry.

To further explore connections between *S. cerevisiae* gene expression and region or vineyard site, we identified genes that were predictive for a specific region or site across all models (local importance, see methods). Only one gene was important across all models for predicting the site OR1 (VIT_0003506001; *V. vinifera* pathogenesis-related protein 10.3). This suggests that we have limited resolution into the specific gene expression patterns that differentiate individual sites using this method. Given that gene expression is inherently noisy (65), increasing observations per vineyard site may improve accuracy and inference from site-specific models in the future.

In contrast to site-specific models, an average of 22.4 genes per region (SD = 13.5) were predictive across all models, with an average of 14.4 genes (SD = 8.4) expressed by *S. cerevisiae* (**Table S7**). Interestingly, many genes that were important for predicting one region were also important for predicting other regions (*BET2, BET3, BIO4, EXG2, FAS2, HEM12, LOH1, MEP3, MRX21, NPT1, PSA1, SNZ3, THI11, THI13, THI72, TUB4*), suggesting that expression of these genes differed consistently between regions. These genes encode proteins involved in diverse cellular processes, including heme biosynthesis, cell wall assembly, and synthesis and transport of fatty acids and nitrogen-containing compounds. While the underlying biochemical processes that lead to consistent expression of these genes within regions remains unknown, we investigated whether initial nitrogen content in grape must was related to the importance of *MEP3*, a gene that encodes an ammonia permease, in predicting a region. Interestingly, *MEP3* was important for predicting the three regions with the lowest average initial yeast assimilable nitrogen (OR, AV, RRV) as well as the region with the second highest yeast assimilable nitrogen (SMV) across vintages (**Figure S7**). Given that nitrogen availability plays a fundamental role in shaping fermentations (66), this relationship was expected. We also noted that four genes associated with thiamine availability were important for predicting multiple regions. This suggests that thiamine availability may drive regional differences in wine outcomes, a postulate that could be measured in a future vintage.

Taken together, these results identify genes directly linked to wine sensory and chemistry that are strong indicators of vineyard region in Pinot noir fermentations. These findings provide a concrete starting point for future investigation into vineyard specific factors that are responsible for wine fermentation outcomes and wine sensory characteristics.

## Conclusion

Microbial biogeography of wine has been documented in globally distributed appellations (4, 7, 8, 14–21), and has been correlated with wine fermentation outcomes (15, 22). In inoculated co-cultures, non-*Saccharomyces* microorganisms both contribute to fermentation and change the behavior of the dominant fermenter *S. cerevisiae*, leading to measurable differences in wine aroma and composition (36–38). Here, we show that grape must ribosomal DNA profiles do not correlate with detected eukaryotic gene expression patterns during primary fermentation. Given that we detected little to no correlation between fungal profiles in initial grape must and genes expressed by those organisms during primary fermentation, DNA profiles may not be a robust indicator for inferring contributions from these organisms in wine sensory outcomes in inoculated fermentations. However, DNA profiles, in particular bacterial profiles, are predictive of vineyard site and retain signatures of site-specific processes such as total precipitation during the growing season. These profiles are rich indicators of the patterns that shape the microbial ecology of grapes, and reflect differences among vineyard sites and regions, even when the same clone (e.g., *Vitis vinifera* L. cv. Pinot noir clone 667) is grown on each site.

In contrast, the gene expression profiles of *S. cerevisiae* and other detected organisms, retain signatures of vineyard site and region as well as the metabolic transformations that occur during fermentation. Using *S. cerevisiae* gene expression as a biosensor for differences between fermentations, we detected site and region specific signatures linked to nitrogen, sulfur, and thiamine metabolism. While these factors are associated with vineyard-specific differences in gene expression profiles, few vineyard site and initial grape must chemical parameters correlate with the transcriptome, which suggests there are still many variables to discover that underlie the complex metabolic activities and gene expression patterns *S. cerevisiae* displays throughout fermentation. In the future, more comprehensive sequencing approaches (e.g., deeper sequencing with methods that capture the full transcriptome, more samples per site) aimed at the factors and organisms identified in this work would allow for a better understanding of these systems. This will need to be accompanied by measurements of many more vineyard, must, and wine characteristics to provide further predictive power and insights into the complexities and subtleties of vineyard specific wine fermentation outcomes.

## Methods

### Grape preparation and fermentation

The winemaking protocol has been described previously (48, 49), but the relevant parts are reproduced with some added details below. The grapes used in this study originated from 15 vineyards in eight American Viticultural Areas in California and Oregon, U.S.A. All grapes were Vitis vinifera L. cv. Pinot noir clone 667, with either rootstock 101-14 (AV1, RRV1, SNC1, SNC2, CRN1, AS1, AS2, SMV1, SMV2, SRH1), Riparia Gloire (OR1, OR2), or 3309C (AV2, RRV2, RRV3). Grapes were hand-harvested grapes at approximately 24 Brix and transported to the University of California, Davis Teaching & Research Winery for fermentation. Grapes were separated into half-ton macrobins on harvest day and Inodose SO_2_ was added to 40 ppm. Upon delivery to the winery, bins were stored at 14°C until the fruit was destemmed and divided into temperature jacket-controlled tanks. N2 sparging of the tank headspace was performed prior to fermentation and tanks sealed with a rubber gasket. We cold soaked the must at 7°C for three days and adjusted TSO_2_ to 40 ppm on the second day. After three days, the must temperature was increased to 21°C and programmed pump overs were used to hold the tank at a constant temperature. Grape must microbiome samples were taken just prior to the increase in temperature. For inoculation, *S. cerevisiae* RC212 was rehydrated with Superstart Rouge at 20 g/hL and inoculated in the must at 25 g/hL. At approximately 24 hours after inoculation, nitrogen content in the fermentations was adjusted using DAP (target YAN - 35 mg/L - initial YAN)/2), and Nutristart (25 g/hL). Nitrogen was adjusted only if YAN was below 250 mg/L. Approximately 48 hours after fermentation, fermentation temperatures were permitted to increase to 27°C, and again added DAP using the formula (target YAN - 35 mg/L - initial YAN)/2, and fermentation were then continued until Brix < 0. Fermentation samples were taken for Brix measurements every twelve hours relative to inoculation and with RNA samples at 2 hours, 6 hours (2019 vintage), 16 hours, 64 hours, and 112 hours (2017 and 2019 vintage). To ensure uniform sampling, a pumpover was performed ten minutes prior to sampling each tank. For RNA samples, 12mL of juice was obtained, centrifuged at 4000 RPM for 5 minutes, supernatant was discarded, and the cell pellet snap frozen in liquid nitrogen. Samples were stored at −80°C until RNA extraction.

### Amplicon sequencing data processing

DNA was extracted for amplicon sequencing and library preparation following (50) and (67). The UC Davis DNA Tech Core performed sequencing using Illumina MiSeq, producing 251 base pair paired-end sequences. We demultiplexed and adapter trimmed libraries by barcode sequences using cutadapt (68). Taxonomically annotated amplicon sequence variant (ASV) counts were generated using DADA2 with the Silva NR database (version 138) for 16S sequences and the UNITE general FASTA release (version 8.2) for ITS sequences (69). All ASVs annotated as “Bacteria,Cyanobacteria,Cyanobacteriia,Chloroplast” and “Bacteria,Proteobacteria,Alphaproteobacteria,Rickettsiales,Mitochondria” were removed as these represent plant mitochondria and chloroplast 16S sequences and not bacterial sequences.

### RNA sequencing data processing

Yeast pellets were thawed on ice, resuspended in 5ml Nanopure water, centrifuged at 2000g for 5min, and aspirated the supernatant. RNA was extracted using the Quick RNA Fungal/Bacterial Miniprep kit including DNAsel column treatment (cat#R2014, Zymo Research). RNA was eluted in 30μL of molecular grade water and assessed for concentration and quality via Nanodrop and RNA gel electrophoresis. Sample concentrations were adjusted to 200ng/μl and used for sequencing. 3’ Tag-seq single-end sequencing (Lexogen QuantSeq) was applied in both the 2017 and 2019 vintage, with the addition of UMI barcodes in 2019. The University of California, Davis DNA Technologies Core performed all library preparation and sequencing.

The first 12 base pairs from each read were hard trimmed and Illumina TruSeq adapters and poly(A) tails were removed. Sourmash gather was used to determine the organisms present in each sample using parameters -k 31 and --scaled 2000 (70, 71). The GenBank microbial database (https://sourmash-databases.s3-us-west-2.amazonaws.com/zip/genbank-k31.sbt.zip) and eukaryotic RNA database (https://osf.io/qk5th/) was used for these queries.

Using results from sourmash, a set of reference genomes was constructed that was representative of all organisms detected within the samples. When different strains of the same species were detected, the one species detected in the largest number of samples was used as a representative species to reduce multi-mapping conflicts. Species present in more than two samples were included because species present in fewer than three samples would have limited predictive power. Species of genus *Saccharomyces* other than *S. cerevisiae* S288C were removed to reduce multi-mapping conflicts. Selected genomes were downloaded from NCBI GenBank; however, if no GTF annotation file was available for the species, the genome and GFF3 file was taken from JGI Mycocosm (72), and the GFF3 was converted to GTF using the R package rtracklayer (73). When no annotation file was available on GenBank or JGI Mycocosm, the genome of the closest species-level strain with a GTF annotation file was used. To find closely related organisms, NCBI taxonomy was searched, selected assemblies were downloaded, and sourmash compare was used with a k-size of 31 (70, 71). The organisms with the highest Jaccard similarity were considered the most similar. When no annotation file was available for similar organisms, an annotation file was generated using WebAugustus (74). See **Table S8** for a description of the best matched genome, the genome used for count generation, and the source of genome annotations. Reference genome FASTA files and GTF files were concatenated together to generate a single reference. STAR was then used to align reads against the constructed reference with parameters --outFilterType BySJout, --outFilterMultimapNmax 20, -- alignSJoverhangMin 8, --alignSJDBoverhangMin 1, --outFilterMismatchNmax 999, -- outFilterMismatchNoverLmax 0.6, --alignIntronMin 20, --alignIntronMax 1000000, -- alignMatesGapMax 1000000, --outSAMattributes NH HI NM MD --outSAMtype BAM, SortedByCoordinate (75). For the 2019 vintage, UMI tools was used to deduplicate alignments (76). The number of reads mapping to each gene was quantified using htseq count using the constructed reference GTF file to delineate gene regions (77).

### RC212 genome assembly and comparison

The *S. cerevisiae* RC212 genome was assembled to estimate the fraction of RNA-sequencing reads in each fermentation originating from non-RC212 *S. cerevisiae* strains. FASTQ files for accession SRR2967888 were downloaded from the European Nucleotide Archive (78). Reads were k-mer trimmed using the khmer trim-low-abund.py command with parameter -k 20 (79) and the Megahit assembler was used with default parameters to assemble reads (80).

### Estimation of non-inoculated yeast in RNA-seq samples

Sourmash gather was used to estimate the fraction of RNA seq reads (k-mers) originating from non-inoculated *S. cerevisiae*. Sourmash gather estimates shared sequence similarity by comparing scaled MinHash signatures derived from k-mer profiles (70, 71). The sourmash Eukaryotic RNA database (https://osf.io/qk5th/) was used, which includes all annotated *S. cerevisiae* genomes in GenBank (e.g., genomes that include *rna_from_genome.fna annotations), as well as our *S. cerevisiae* RC212 genomes assembly.

### Correlation between ribosomal DNA amplicon sequencing data and 3’ Tag-seq data for non-*Saccharomyces* organisms

Fermentations with fungal ITS amplicon sequencing data and 3’ Tag-seq were compared. First, ribosomal DNA amplicon sequencing read counts from *H. uvarum* were regressed against total 3’ Tag-seq counts from *H. uvarum* using counts from 16 hours, 64 hours, and 112 hours of fermentation. 3’ Tag-seq counts were derived from STAR and htseq (see RNA sequencing data processing above). Counts were transformed into compositional counts (relative abundance) prior to linear regression (81). Linear regression was performed using the lm() function in R. This analysis was performed again separately for *H. uvarum* and *A. pullulans* using counts from the 2 hour and 6 hours samples taken in the 2019 vintage. Given that this analysis relied on reads aligned to annotated 3’ regions, a separate regression was performed a using proportion of reads assigned to a given organism derived from sourmash gather (see RNA sequencing data processing above). Only results from the first analysis were reported as R^2^ values were within 0.01 between both analyses.

### ANOSIM and NMDS

Compositional data analysis was used for amplicon and transcriptome counts (81). The transform() function in the microbiome bioconductor package was used to transform counts by centered log ratio (82, 83). To test for differences among groups, Aitchinson distance (Euclidean distance on CLR-transformed counts) was used and tested with the anosim() function in the vegan package using parameters distance = "euclidean" and permutations = 9999 (84, 85). A cut off of p = 0.05 was used for statistical significance. To construct NMDS plots, Aitchinson distance was taken using the metaMDS() function in the vegan package with parameter distance = "euclidean". Results were plotted using the ggplot2 package (86).

### Amplicon sequencing random forest models

Random forest classifiers were built using the R ranger package (87). Using ASV counts produced by DADA2, counts were normalized by dividing by total number of aligned reads. The tuneRanger() function was used in the tuneRanger package to optimize each model for parameters m.try, sample.fraction, and min.node.size (88). The ranger() function was then used to build each model with parameters from tuneRanger as well as num.trees = 10000, importance = "permutation", and local.importance = TRUE. As a supervised technique, random forest classifiers are trained on a subset of data and tested on a separate subset to calculate predictive accuracy. For models built with samples from all vintages, the createDataPartitionQ function in the R caret package was used to randomly but equally partition training and testing sets with a 70:30 split, ensuring that all class labels were equally represented in both sets (89). For other models, the classifier was built using all samples from two vintages and validated on the held-out vintage. Accuracy and kappa statistics were calculated for each model.

### RNA sequencing random forest model

Counts were imported into R and normalized by dividing by total number of aligned reads (e.g., library size). Given that random forests expects independent samples and RNA-sequencing was conducted in time series over the course of primary fermentation, each gene from each time series set was summarized into mean count, minimum count, maximum count, total count, and standard deviation of counts. Variable selection was performed using the vita method (90) and models were built using the same methods as with amplicon sequencing models.

To estimate vineyard-and region-specific gene importance, variable selection and model optimization were performed with 100 different seeds. For each model, gene local importance was averaged for each fermentation from a vineyard site or region in the training set and genes with positive average local permutation importance were retained. The intersection of genes from all models was then taken to determine which genes were predictive for a particular site or region in all models. Although random forests were trained on summarized gene attributes, any genes that were predictive across any attribute were retained as these attributes were often highly correlated.

### Mantel tests

Mantel tests were performed to assess the similarity between samples across measurements of bacterial abundance, fungal abundance, transcriptome abundance, initial grape must chemistry, and vineyard site parameters (91, 92). The Mantel test determines the correlation between the same samples in different matrices, testing whether similarities between samples estimated from one measurement type match similarities of the same samples estimated from a different measurement type (91, 92). These tests were performed using complete cases, with microbiome and transcriptome abundances from the 2017 and 2019 vintages. Vineyard site parameters total precipitation and growing degree days were estimated using the PRISM climate models including dates April 1 - September 30 in 2017 and 2019 (93). Distance matrices were calculated for each matrix using the dist() function in R, with parameters method = "euclidean", with the exception of geographic distance which was calculated using the distm() function in the package geosphere with parameter distHaversine (94). When distances for disparate measurement types were calculated at the same time, values were first scaled and centered using the function scale() with parameters center = TRUE and scale = TRUE. Mantel tests were performed with the mantel() function in the vegan package with parameters method = "spearman", permutations = 9999, and na.rm = TRUE (85, 92). p value adjustment were applied using the function p.adjust() with parameter method = "fdr" and a false discovery rate of p = 0.1 used.

## Data Availability

RNA sequencing data is available in the Sequence Read Archive under accession number PRJNA680606. Microbiome data is available under accession numbers PRJNA642839 and PRJNA682452. All analysis code is available at github.com/montpetitlab/Reiter_et_al_2020_SigofSite.

## Acknowledgements

We thank all past and current members of the Steenwerth, Runnebaum, and Montpetit laboratories for their support of this work, as well as the students and staff of the UC Davis Pilot Winery. T.R. was supported by the Gordon and Betty Moore Foundation's Data-Driven Discovery Initiative [GBMF4551]; Harry Baccigaluppi Fellowship; Horace O Lanza Scholarship; Louis R Gomberg Fellowship; Margrit Mondavi Fellowship; Haskell F Norman Wine & Food Fellowship; Chaîne des Rôtisseurs Scholarship; Carpenter Memorial Fellowship. The authors would like to recognize support from Jackson Family Wines, in addition to support from Lallemand Inc.

---

**Figure S1:**
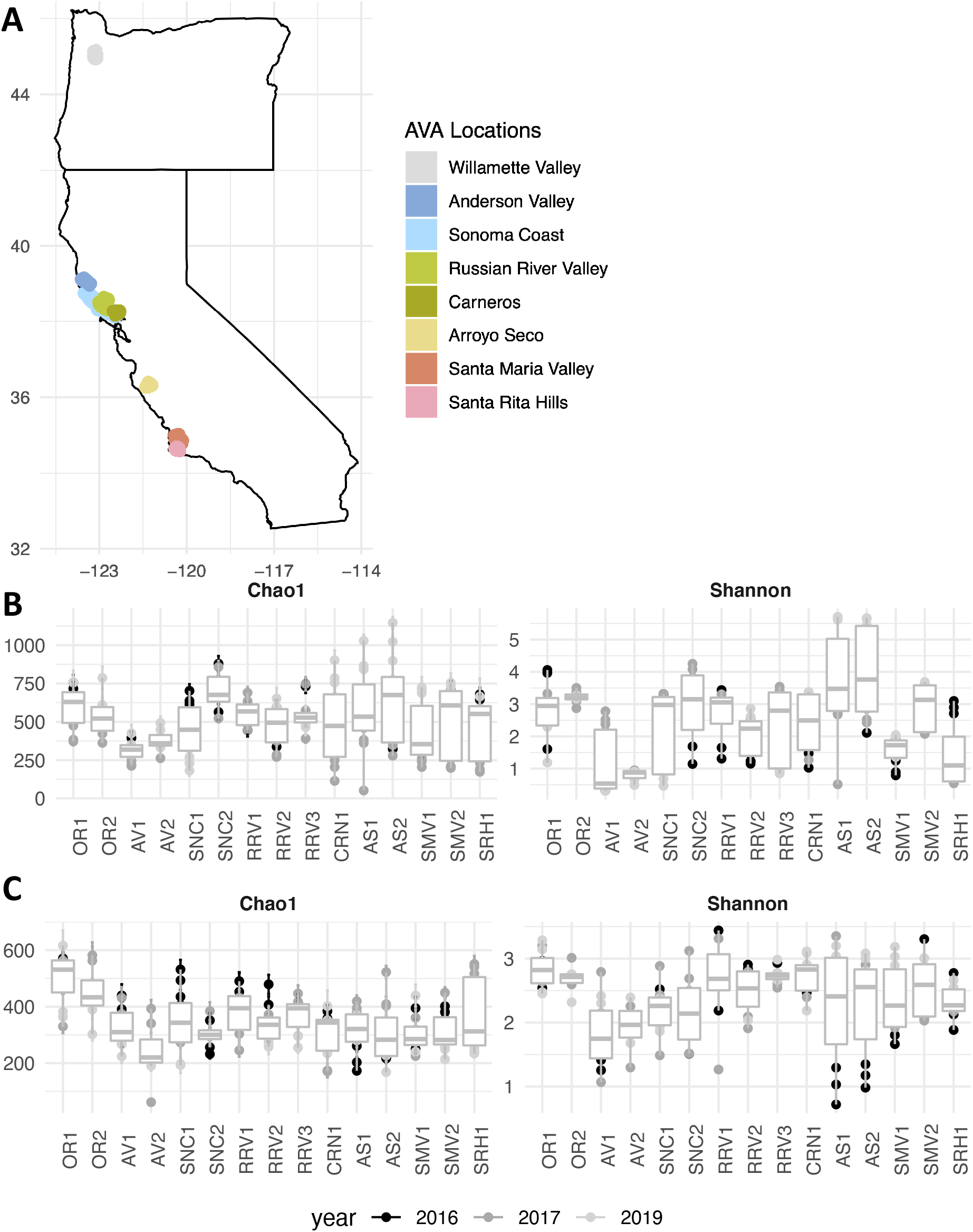
Diversity of vineyards and ribosomal DNA profiles in this study. **A)** Map displaying the 15 vineyard locations across eight American Viticultural Areas (AVAs) in California and Oregon. **B, C)** Bacterial and fungal ribosomal DNA amplicon sequencing Chao 1 and Shannon alpha diversity for mean species diversity per vineyard site, averaged across vintages. **B** Bacteria. **C** Fungi.

**Figure S2:**
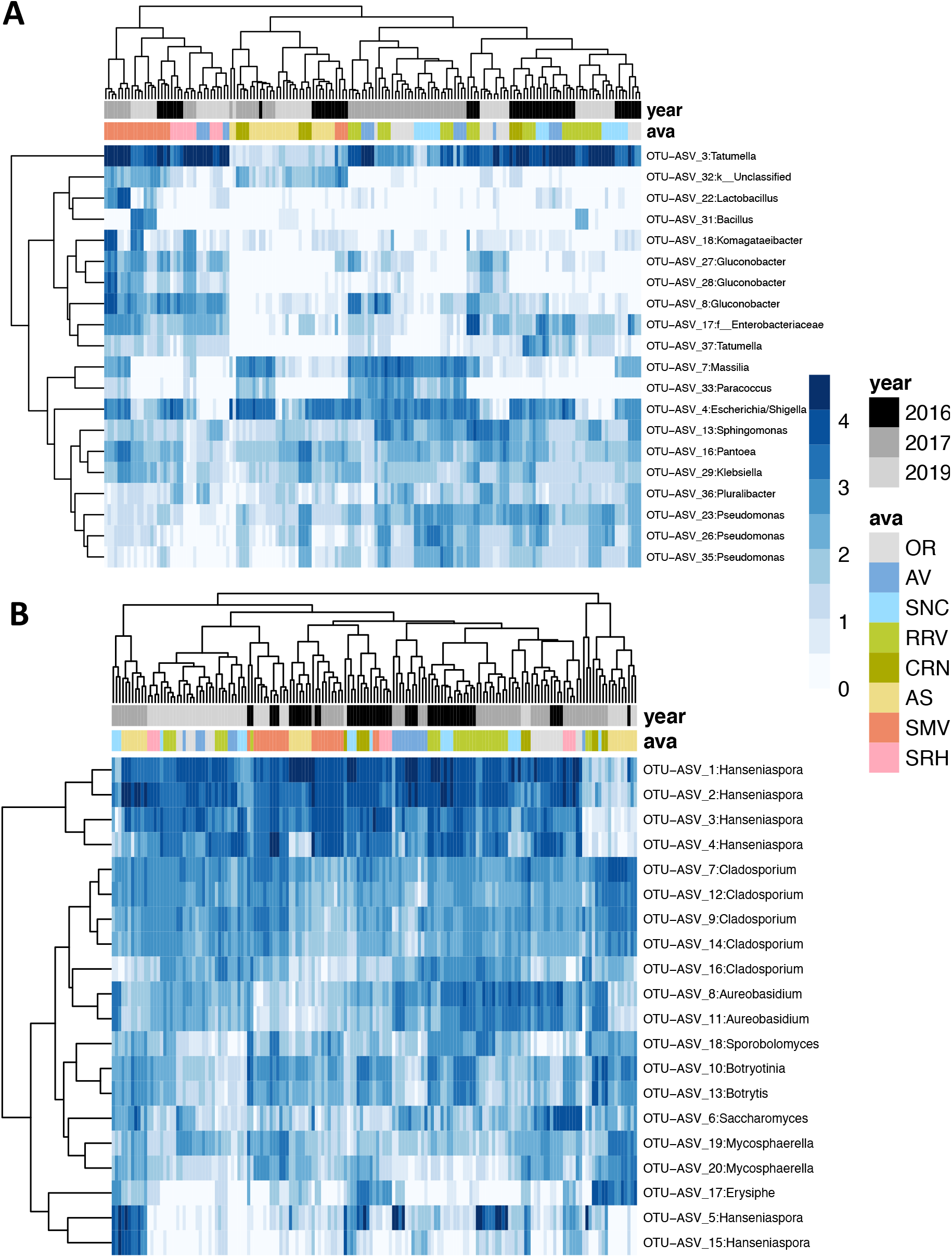
Some ribosomal sequencing variants were detected across vineyards and vintages. *Top* 20 most abundant ribosomal DNA amplicon sequencing variants across vintages. Labelled as genus or the next lowest taxonomic rank of classification. **A** Bacteria. **B** Fungi. *Tatumella* was the most abundant bacterial amplicon sequencing variant across vineyards and vintages, while *Hanseniaspora* was the most abundant fungal amplicon sequencing variant.

**Figure S3:**
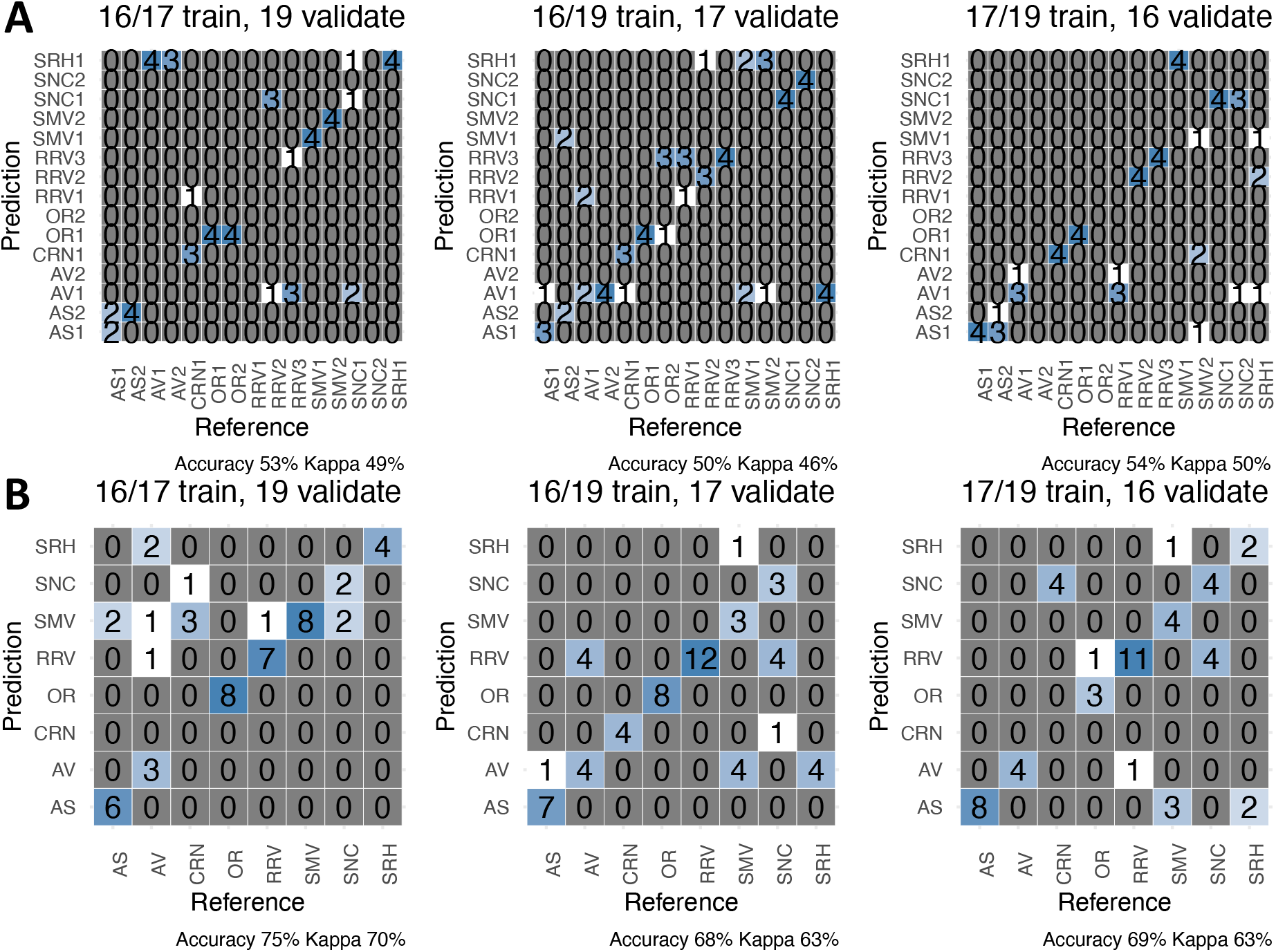
Accuracy of random forests models using bacterial ribosomal DNA profiles. Confusion matrices depicting accuracy of random forests models built with bacterial ribosomal DNA amplicon sequencing data to predict **A)** vineyard site and **B)** vineyard region. The models depicted were trained on two vintages and validated on the third.

**Figure S4:**
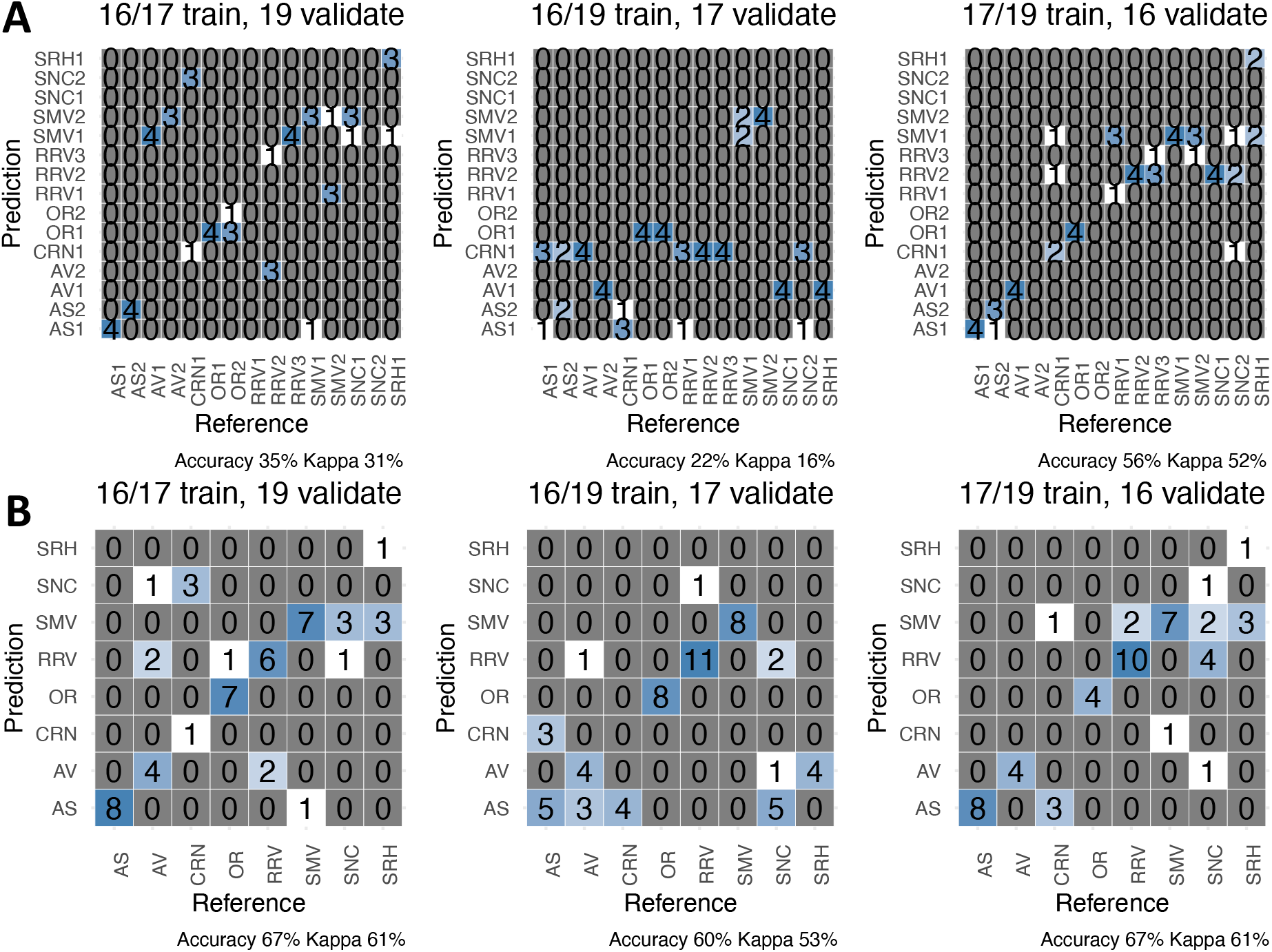
Accuracy of random forests models using fungal ribosomal DNA profiles. Confusion matrices depicting accuracy of random forests models built with fungal ribosomal DNA amplicon sequencing data to predict **A)** vineyard site and **B)** vineyard region. The models depicted were trained on two vintages and validated on the third.

**Figure S5:**
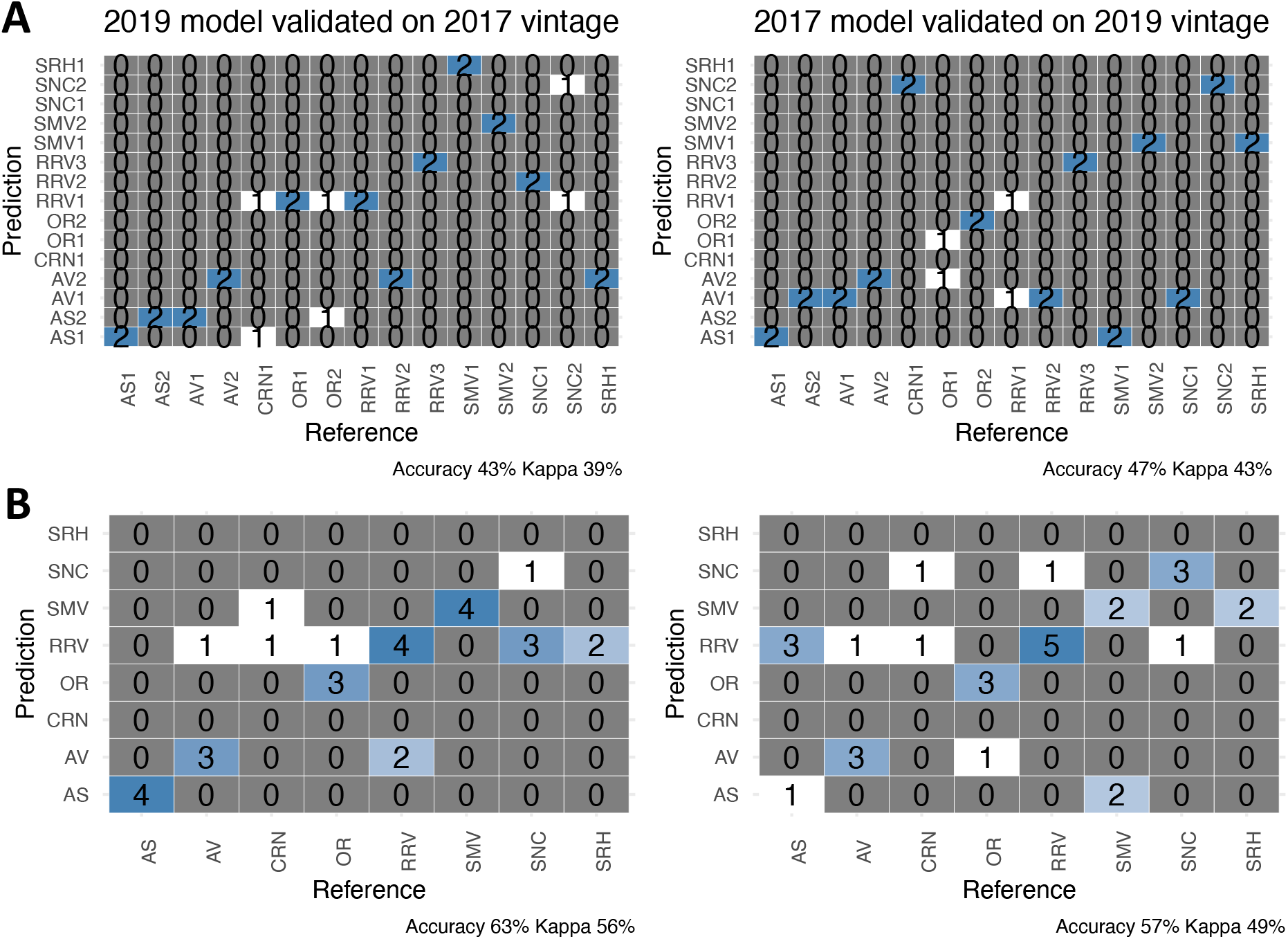
Accuracy of random forests models using RNA sequencing. Confusion matrices depicting accuracy of random forests models built with RNA sequencing data to predict **A)** vineyard site and **B)** vineyard region. The models depicted were trained on one vintage and validated on the other.

**Figure S6:**
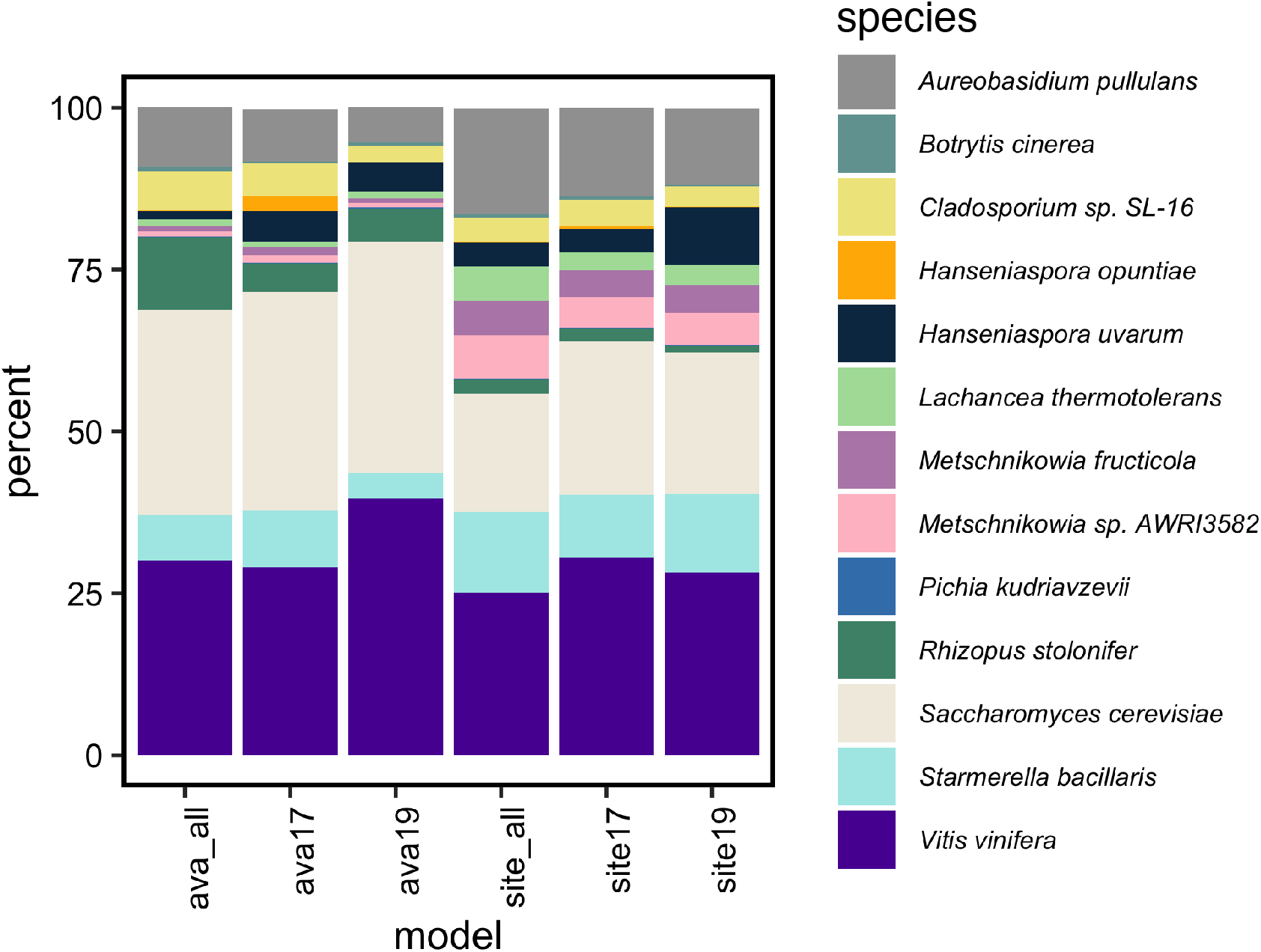
Percent of accuracy attributable to different organisms in random forests models. Importance of genes expressed by different organisms in the overall model. A higher percentage of variable importance was attributable to *S. cerevisiae* and *V. vinifera* in models that predicted region than vineyard site.

**Figure S7:**
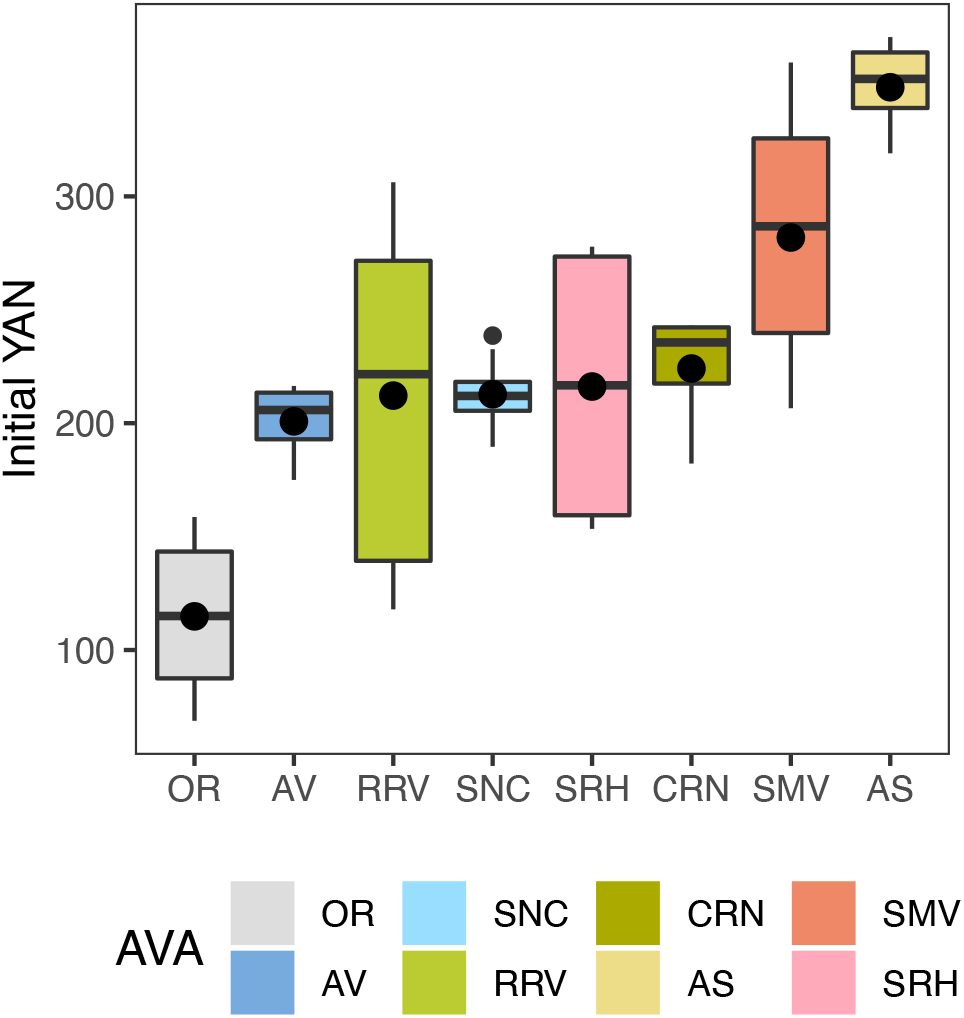
Initial yeast assimilable nitrogen (YAN) in grape musts across vintages. Black dots mark the mean initial YAN value calculated from all fermentations in the 2017 and 2019 vintages. *MEP3*, which encodes an ammonia permease, was important for predicting the three regions with the lowest average initial YAN (OR, AV, RRV) and the region with the second highest initial YAN (SMV).

## Supplemental Data Tables

For Tables S2 - S7: see accompanying supplemental files.

**Table S1:** Accuracy of random forests models built with fungal and bacterial ribosomal DNA amplicon sequencing data and transcriptome data. Validation set “30” indicates models that were trained with 70% of data and validated on the held-out 30%.

| Data set | Model type | Validation set | Accuracy |
| --- | --- | --- | --- |
| bacteria | AVA | 2019 | 0.74 |
| bacteria | AVA | 2017 | 0.68 |
| bacteria | AVA | 2016 | 0.69 |
| bacteria | AVA | 30 | 0.93 |
| bacteria | site | 2019 | 0.53 |
| bacteria | site | 2017 | 0.5 |
| bacteria | site | 2016 | 0.54 |
| bacteria | site | 30 | 0.93 |
| fungi | AVA | 2019 | 0.67 |
| fungi | AVA | 2017 | 0.6 |
| fungi | AVA | 2016 | 0.67 |
| fungi | AVA | 30 | 0.95 |
| fungi | site | 2019 | 0.33 |
| fungi | site | 2017 | 0.23 |
| fungi | site | 2016 | 0.56 |
| fungi | site | 30 | 0.89 |
| transcriptome | AVA | 2019 | 0.57 |
| transcriptome | AVA | 2017 | 0.63 |
| transcriptome | AVA | 30 | 0.87 |
| transcriptome | site | 2019 | 0.43 |
| transcriptome | site | 2017 | 0.43 |
| transcriptome | site | 30 | 0.67 |

**Table S8:**
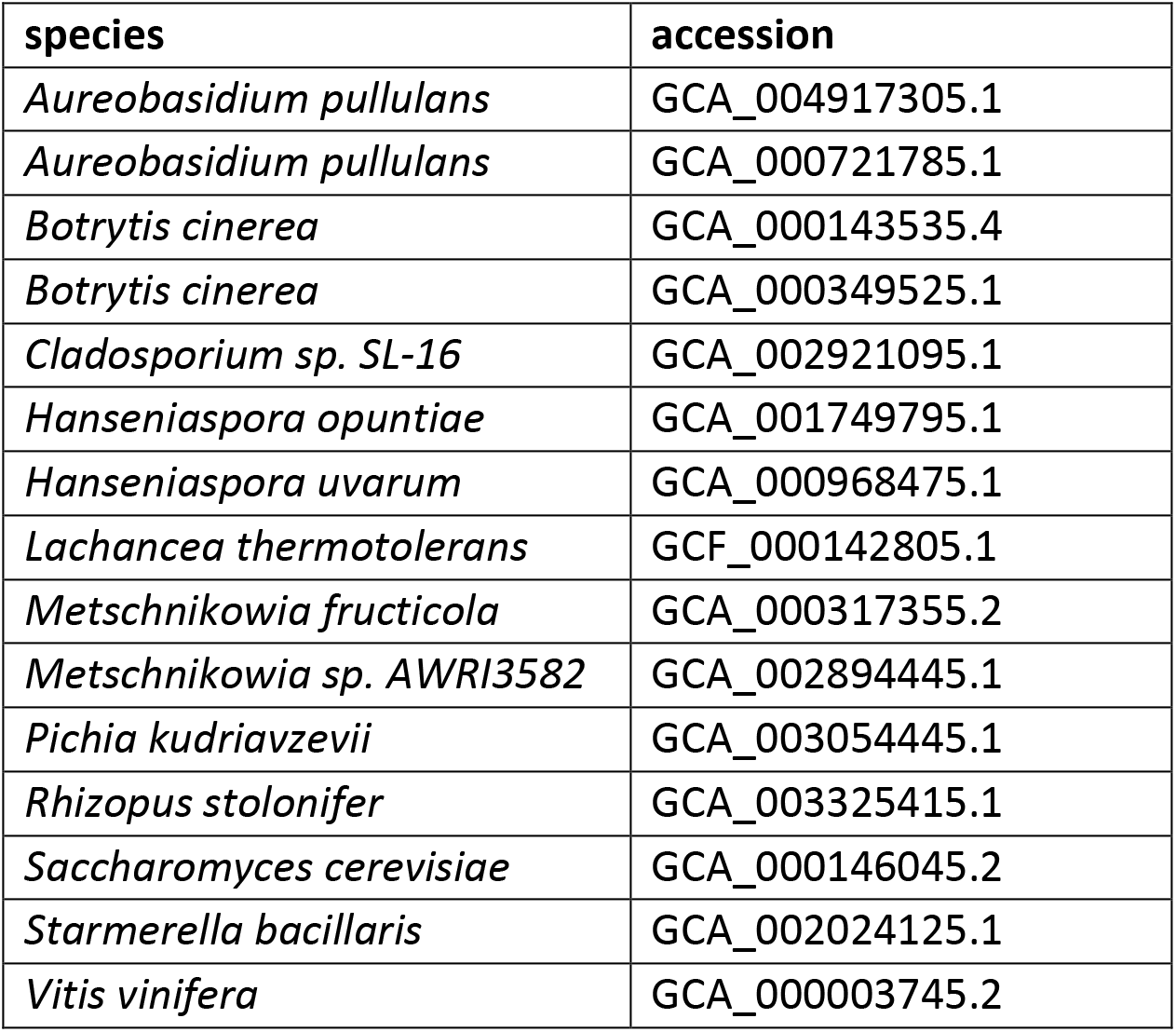
Species names and accession numbers for genomes used in this study.

| species | accession |
| --- | --- |
| *Aureobasidium pullulans* | GCA_004917305.1 |
| *Aureobasidium pullulans* | GCA_000721785.1 |
| *Botrytis cinerea* | GCA_000143535.4 |
| *Botrytis cinerea* | GCA_000349525.1 |
| *Cladosporium sp. SL-16* | GCA_002921095.1 |
| *Hanseniaspora opuntiae* | GCA_001749795.1 |
| *Hanseniaspora uvarum* | GCA_000968475.1 |
| *Lachancea thermotolerans* | GCF_000142805.1 |
| *Metschnikowia fructicola* | GCA_000317355.2 |
| *Metschnikowia sp. AWRI3582* | GCA_002894445.1 |
| *Pichia kudriavzevii* | GCA_003054445.1 |
| *Rhizopus stolonifer* | GCA_003325415.1 |
| *Saccharomyces cerevisiae* | GCA_000146045.2 |
| *Starmerella bacillaris* | GCA_002024125.1 |
| *Vitis vinifera* | GCA_000003745.2 |

## Notes

https://github.com/montpetitlab/Reiter_et_al_2020_SigofSite

### Competing Interest Statement

The authors have declared no competing interest.

### Summary of Updates

Supplemental Tables added.

